# SEX DIFFERENCES IN μ-OPIOID REGULATION OF COERULEAR-CORTICAL TRANSMISSION

**DOI:** 10.1101/2020.10.12.336552

**Authors:** Herminio M Guajardo, Rita J Valentino

## Abstract

Stress-induced activation of locus coeruleus (LC)-norepinephrine (NE) projections to the prefrontal cortex is thought to promote cognitive responses to stressors. LC activation by stressors is modulated by endogenous opioids that serve to restrain LC activation and to facilitate a return to baseline activity upon stress termination. Sex differences in this opioid influence could be a basis for sex differences in stress vulnerability. Consistent with this, we recently demonstrated that μ-opioid receptor (MOR) expression is decreased in the female rat LC compared to the male LC and this was associated with sexually distinct consequences of activating MOR in the LC on cognitive flexibility. Given that the LC-NE system affects cognitive flexibility through its projections to the medial prefrontal cortex (mPFC), the present study quantified and compared the effects of LC-MOR activation on mPFC neural activity in male and female rats. Local field potential (LFPs) were recorded from the mPFC of freely behaving male and female rats before and following local LC microinjection of the MOR agonist, DAMGO or vehicle. Intra-LC DAMGO altered the LFP power spectrum selectively in male, but not female rats, resulting in a time-dependent increase in the power in delta and alpha frequency bands. LC microinfusion of ACSF had no effect in either sex. Together, the results are consistent with previous evidence for decreased MOR function in the female rat LC and demonstrate that this translates to a diminished effect on cortical activity that can account for sex differences in cognitive consequences. Decreased LC-MOR function in females could contribute to greater stress-induced activation of the LC, and increased vulnerability of females to hyperarousal symptoms of stress-related neuropsychiatric pathologies.

## INTRODUCTION

Stress-related neuropsychiatric pathologies are more prevalent in females relative to males. For example, post-traumatic stress disorder (PTSD), and depression are nearly two times more prevalent in females when compared with males (Kessler *et al*, 1994; Kessler *et al*, 1995). As these neuropsychiatric diseases are often characterized by symptoms of hyperarousal, sex differences in brain arousal systems and their regulation by stress may contribute to the female bias their prevalence (Wong *et al*, 2000).

One neurotransmitter system that is involved in stress and arousal and that exhibits sex differences is the locus coeruleus (LC)-norepinephrine (NE) system. LC neuronal discharge rate is positively correlated with arousal state (Aston-Jones and Bloom, 1981; Berridge and Foote, 1991; Berridge *et al*, 1993) and LC hyperactivity is thought to underlie the core feature of hyperarousal that characterizes stress-related neuropsychiatric diseases (Gold and Chrousos, 1999; Gold *et al*, 1996; Koob, 1999; O’Donnell *et al*, 2004). In response to an acute stressor, LC neurons are co-regulated in an opposing manner by the stress-related neuropeptide, corticotropin-releasing factor (CRF), which activates LC neurons and the endogenous opioid, enkephalin, which inhibits LC neurons (Curtis *et al*, 2001; Curtis *et al*, 2012). The CRF-induced excitation of LC neurons predominates during the acute stress response and this activation is integral to stress-induced arousal (Page *et al*, 1993). The CRF-elicited LC activation during stress may also promote cognitive flexibility because CRF microinfusion into the LC in doses that produce the same magnitude of LC activation as stressors facilitate attentional set shifting and reversal learning in attentional set shifting tasks (Snyder *et al*, 2012; Valentino and Van Bockstaele, 2008). In contrast to CRF, endogenous opioids released in the LC during stress act at μ-opioid receptors (MOR) to temper CRF-mediated activation and promote recovery of LC discharge rate when the stressor is terminated (Curtis *et al*, 2001; Curtis *et al*, 2012).

Sex differences in function of the LC-norepinephrine system arise from differences in sensitivity to both CRF and endogenous opioids. Previous rodent studies provided evidence that the CRF1 receptor (CRF1) exhibits increased Gs-protein coupling and decreased association to β-arrestin 1 in females, resulting in increased sensitivity of LC neurons to CRF and decreased stress- and agonist-induced internalization (Bangasser *et al*, 2010; Bangasser *et al*, 2013). In contrast MOR function is decreased in female LC neurons. For example, MOR mRNA and protein is decreased in female LC compared to males and this is associated with a lower magnitude of MOR-mediated inhibition by the peptide agonist, DAMGO (Guajardo *et al*, 2017). Consistent this, DAMGO microinjection into the LC produces a greater impairment in cognitive flexibility in males compared to females (Guajardo *et al*, 2017).

Because the LC-NE system influences cognitive flexibility through its projections to the medial prefrontal cortex (mPFC), the present study was designed to determine how molecular and cellular sex differences at the level of LC-MOR translate to changes in neural activity in the mPFC. The effect of activating MOR in the LC on mPFC network activity was compared in male and female rats.

## MATERIALS AND METHODS

### Animals

Adult male (n=4) and female (n=4) Sprague Dawley rats (Charles River, Wilmington, MA) were shipped from the vendor at ~70 days of age. Experiments were conducted 1 week after arrival. Rats were singly housed in a climate-controlled room with a 12-h light–dark cycle (lights on at 0700 hours). Food and water were freely available. Female rats were intact. Animal use and care was approved by the institutional animal care and use committee of the Children’s Hospital of Philadelphia.

### Surgery

Male and female rats were implanted with a single cannula guide into the LC (26 gauge, Plastics One, Roanoke, VA). Rats were anesthetized with isofluorane (2%) and positioned in a stereotaxic instrument with the head tilted at a 15° angle to the horizontal plane (nose down). A hole was drilled centered at LC coordinates relative to lambda: AP −3.6 mm, ML ±1.1 mm, and the guide cannula was lowered to 5.0 mm below brain surface. The guide cannula was affixed to skull and skull screws with cranioplastic cement. An obdurator was inserted into the guide cannula to prevent occlusion. During the same surgery, a depth electrode (100 μm) (Microprobes for life science, Gaithersburg, MD) was implanted in the mPFC (+3.2 AP, −0.6 ML, 3.0 DV) ipsilateral to the LC-cannula guide for recording local field potentials (LFP). A ground wire was attached to the skull and skull screws. Animals were allowed 5 days to recover before experimental manipulations.

### Drugs for intra-LC microinjections

Drugs used for intra-LC microinfusion were DAMGO ([D-Ala2, N-MePhe4, Gly-ol]-enkephalin; Abcam, Cambridge, MA), a synthetic opioid peptide with high MOR specificity and clonidine HCl (Sigma, St Louis, MO). These compounds were aliquoted and concentrated using a Speed Vac and stored at −20°C before experimental procedures. On the day of the experiments, drugs were dissolved in artificial cerebrospinal fluid (ACSF). The doses that were microinfused into the LC were DAMGO (10 pg in 200 nl) and clonidine (50 ng in 200 nl). The dose of DAMGO is one that has been demonstrated to produce sexually distinct effects on LC activity and on cognitive endpoints when microinfused into the LC (Guajardo *et al*, 2017). The dose of clonidine has been demonstrated to produce a long lasting cessation of LC spontaneous discharge (Page *et al*, 1993).

### Electrophysiological recordings

Experiments began at least 5 days after surgery. All mPFC recordings were performed in the unanesthetized state. For each rat in the study, cables connected the head-stage to a data acquisition system (AlphaLab; Alpha Omega; Nazareth Israel). Cannula (26 gauge, Plastics One, Roanoke, VA) were inserted into the guides and secured using a connector that screws around the cannula guide when inserted. Tubing connecting the cannula to the syringe pump was threaded through a flexible coil that allowed free movement of the rat yet maintained stability of the cannula in the guide. For all animals in the study, pre-injection LFP recordings were taken for 60 minutes. This allowed the rat to undergo multiple sleep/wake cycles that could be used to validate the mPFC LFP recording. LFP activity was recorded and amplified at a gain of 5000 Hz, bandwidth of 1-350 Hz. Following the pre-injection recording period, ACSF (200 nl), DAMGO (10 pg in 200 nl), or clonidine (50 ng in 200 nl) was microinfused into the LC. Recordings continued for 60 minutes after drug administration. ACSF and DAMGO were administered to the same subjects on different days with treatments being 7 days apart.

### Histology

After the experiment rats were anesthetized with isofluorane, pontamine sky blue dye (PSB, 200 nl) was injected through the LC cannula to verify placement. Brains were removed, and frozen 30 μm-thick coronal sections were cut on a cryostat and mounted on pre-cleaned plus slides. The sections were stained with neutral red dye for localization of the PSB spot (Figure 1A).

**Figure 1.**
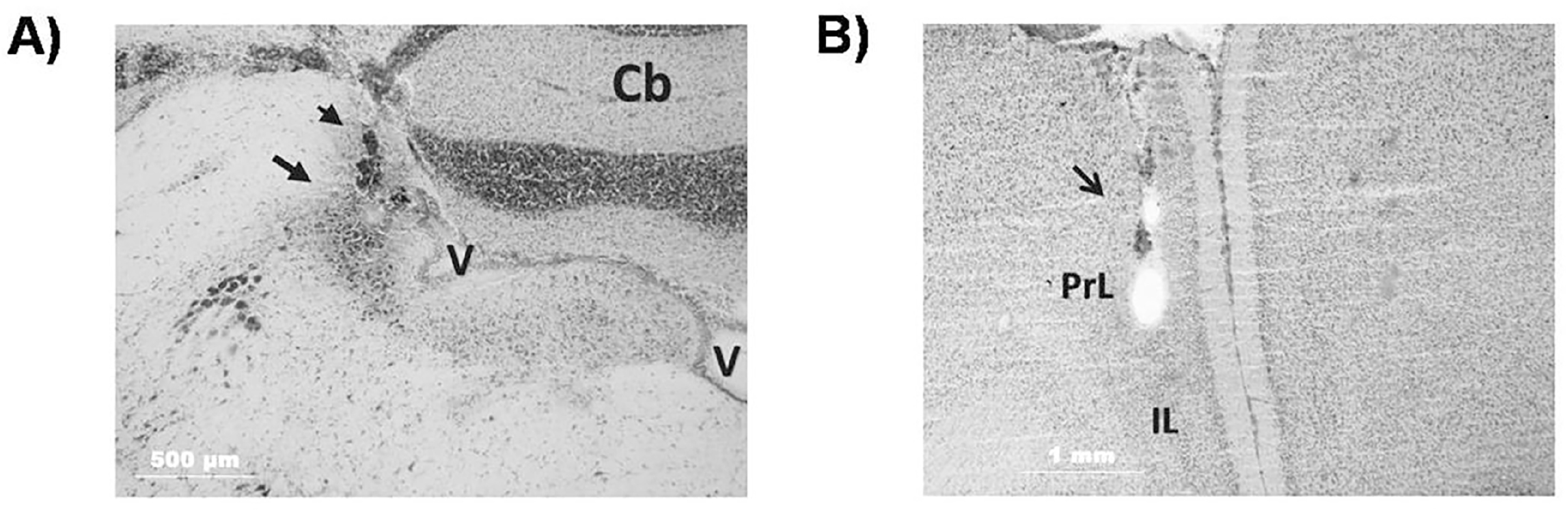
Histological verification of intra-LC injection and mPFC electrode placement. Brightfield photomicrograph of a Neutral red counterstained section through the LC showing histological verification of the single injection site. (A) Arrow points to the LC and the arrowhead points to the dye, which is localized to the LC (Cb, cerebellum; V ventricle). (B) Neutral red counterstained section through the prelimbic mPFC (PrL) showing histological verification of the recording electrode placement. Arrow points to the electrode track (IL, infralimbic mPFC).

### Data Analysis

LFP raw traces were time stamped in Spike2 to remove noise and converted to Power Spectra Density (PSD) raw plots indicating the power in 128 bins from 0 to 20 Hz using Neuroexplorer (Nex Technologies, Madison, AL). The power in different frequency bands (delta, 2-4 Hz; theta, 4-8 Hz; alpha, 8-12 Hz; and beta, 12-20 Hz) was calculated for each rat. A two-way repeated measure analysis of variance (rmANOVA) with sex as the between factor and time with respect to injection as the repeated measure for each individual frequency band. The Tukey’s HSD was used for post-hoc comparisons between means. An alpha level of p<0.05 was the maximum threshold for statistical significance. Post hoc tests were only performed if an interaction is indicated.

## RESULTS

### Effects of intra-LC DAMGO on cortical activity

DAMGO was microinfused into the LC of 4 male and 4 female rats. Figure 1 shows an example of histological verification of the injection site into the LC and the location of injections in all rats. Within 100 s of the local LC injection of DAMGO mPFC synchronization increased primarily in male rats. The representative spectrograms displaying power in different frequencies over time show an increase in amplitude spanning 1-10 Hz that lasts at least 10 min in males and increase in power at 1-2 Hz lasting only about 1-2 min in females. (Figure 2A, B). LFPs were analyzed as PSD plots and the power in different frequency bands at different time blocks were analyzed and compared between sexes (Figure 2C-2D). A two-way repeated measures (rm) ANOVA was done to compare power within individual frequency bands with sex as the between factor and time with respect to injection as the repeated measure for each individual frequency band. Analysis of power in the delta frequency band indicated no effect of Sex (F (1, 6) =3.84, p<0.09), there was an effect of Time (F (3, 4) =21.50, p<0.006), and a Sex X Time interaction (F (3, 6) =28.4, p<0.003). Analysis of power in the theta frequency band indicated a trend for effect of Sex (F (1, 6) =5.34, p=0.06), no effect of Time (F (3, 4) =4.51, p=0.09) and a trend for a Sex X Time interaction for power in the theta frequency band (F (3, 4) =6.2, p=0.05). Analysis of power in the alpha frequency band indicated an effect of Sex (F (1, 6) =11.20, p<0.01), no effect of Time (F (3, 4) =3.77, p=0.1) and a Sex X Time interaction (F (3, 4) =6.93, p<0.04). Analysis of power in the beta frequency band indicated no effect of Sex (F (1, 6) =3.80, p=0.1), no effect of Time (F (3, 4) =4.14, p=0.1) and there was no Sex X Time interaction (F (3, 4) =4.75, p<0.08).

**Figure 2.**
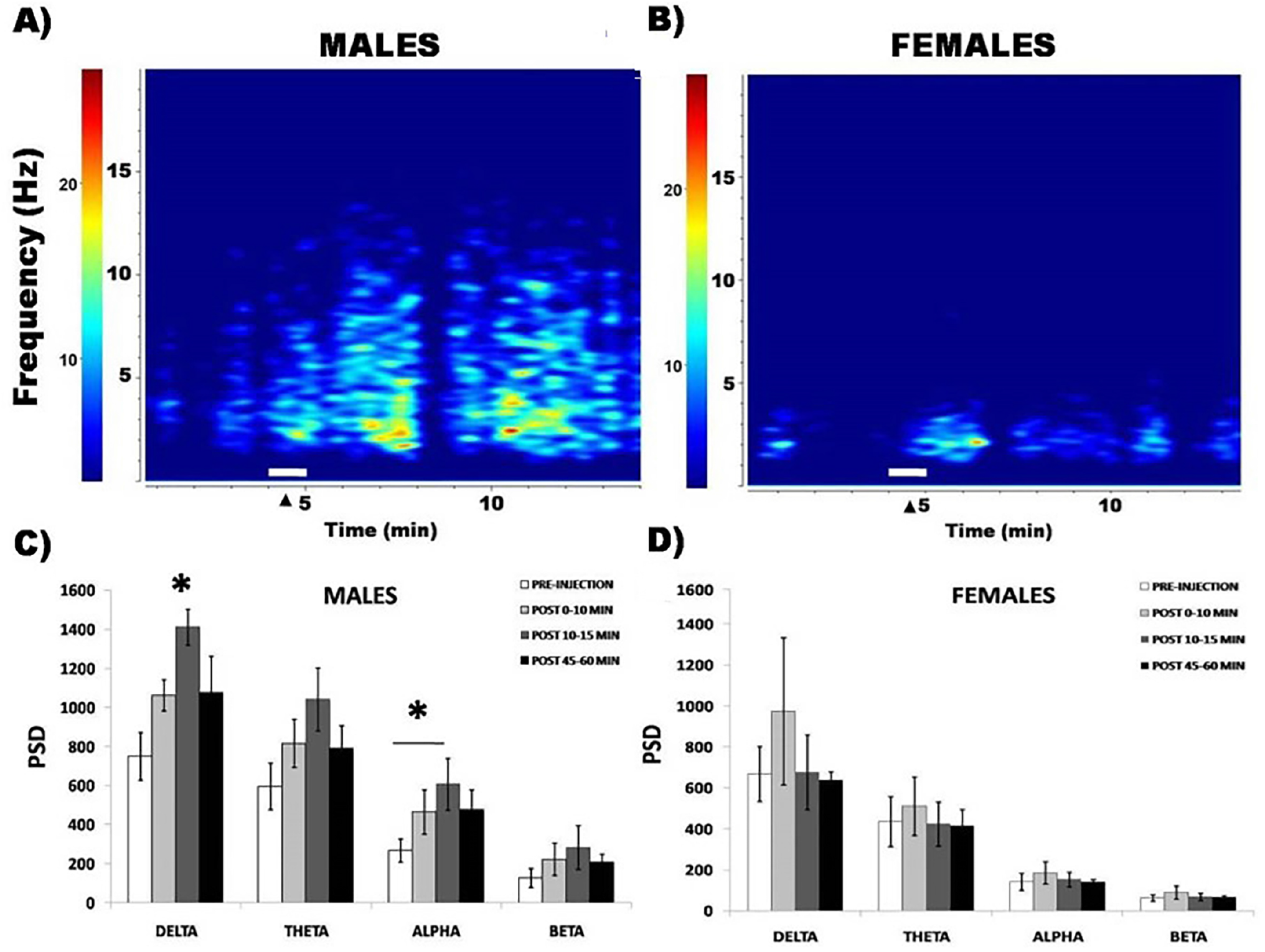
Intra-LC DAMGO resulted in a time-dependent synchronization of mPFC activity in male but not female rats. Representative time-frequency spectrograms of mPFC LFP activity generated from a male and female rat. (A and B) Power in different frequencies (ordinate) is indicated as color (hotter colors being greater). DAMGO (10 pg in 200 nl) was microinfused into the LC at white horizontal line and arrowhead. (C and D) The bar graphs represent the mean raw power spectral density values at different frequency bands (delta 2-4 Hz, theta 4-8 Hz, alpha 8-12 Hz and beta 12-20 Hz) in male and female rats, respectively, determined 10 min before (pre-injection), 0-10 min, 10-15 min, and 45-60 min after intra-LC DAMGO on mPFC network activity. Error bars represent ± SEM. *p<0.05.

The post-hoc comparisons between means within specific frequency bands at different time blocks after DAMGO administration (0 to 10 minutes, 10 to 15 minutes, and 45 to 60 minutes) revealed a time-dependent synchronization of mPFC activity such that there was no significant effect of DAMGO in any frequency band at 0-10 minutes after injection (Figure 2C and 2D). Notably, DAMGO significantly increased power in the delta and alpha frequency bands at 10-15 minutes after administration in males only (delta, p<0.05; alpha, p<0.05, Tukey’s HSD) (Figure 2C and 2D). At this time period, there was also a trend for DAMGO to increase power in the theta frequency band selectively in male rats (p=0.05). DAMGO effects recovered by 45-60 min post-injection.

### Lack of effect of intra-LC ACSF on mPFC network activity

In contrast to DAMGO, local ACSF injection into the LC did not affect mPFC activity of the same rats (Figure 3). A two way ANOVA comparing power in different frequency bands at different post-injection times revealed an effect of Sex (F (1, 6) = 7.5, p=0.03), no effect of Time (F (3, 4) =0.39, p=0.8) and no Sex X Time interaction for power in the delta (F (3, 4) =5.21, p=0.07). Analysis of power in the theta frequency band indicated no effect of Sex (F (1, 6) =3.27, p=0.1), no effect of Time (F (3, 4) =0.85, p=0.53) and there was no Sex X Time interaction (F (3, 4) = 1.39, p=0.4). Analysis of power in the alpha frequency band indicated no effect of Sex (F (1, 6) =1.32, p=0.3), no effect of Time (F (3, 4) =0.41, p=0.8) and there was no Sex X Time interaction (F (3, 4) = 2.61, p=0.2). Analysis of power in the beta frequency band indicated no effect of Sex (F (1, 6) =1.24, p=0.3), no effect of Time (F (3, 4) =0.57, p=0.7) and there was no Sex X Time interaction (F (3, 4) = 3.30, p=0.14).

**Figure 3.**
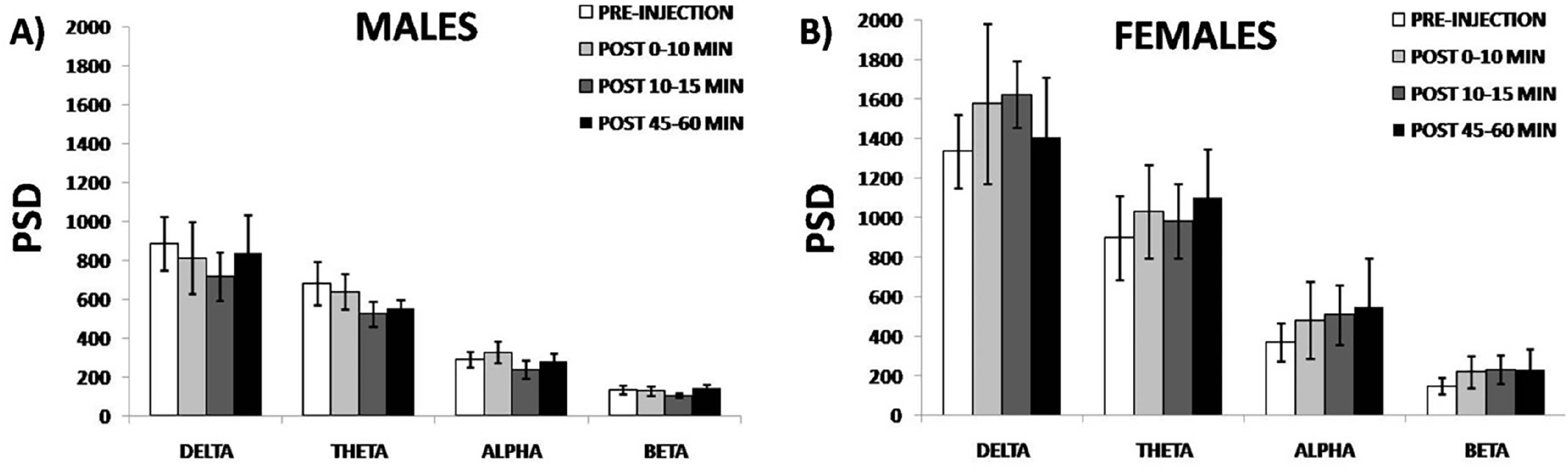
Lack of effect of intra-LC ACSF on mPFC network activity. Bar graphs represent the mean raw power in different frequency bands at different times before and after intra-LC ACSF (200 nl). A) Males and B) Females at different time blocks (0-10, 10-15, 45-60 minutes after Intra-LC DAMGO injection). Error bars represent ± SEM.

### Comparable effects of intra-LC clonidine on cortical activity of male and female rats

Like DAMGO, clonidine inhibits LC discharge rate, and intra-LC injection increases cortical electroencephalographic synchrony of anesthetized rats (Berridge *et al*, 1993). Consistent with this, intra-LC microinfusion of clonidine (50 ng) increased the LFP amplitude in rats not anesthetized. Notably, this effect appeared comparable in a male and female rat (Figure 4A-4D).

**Figure 4.**
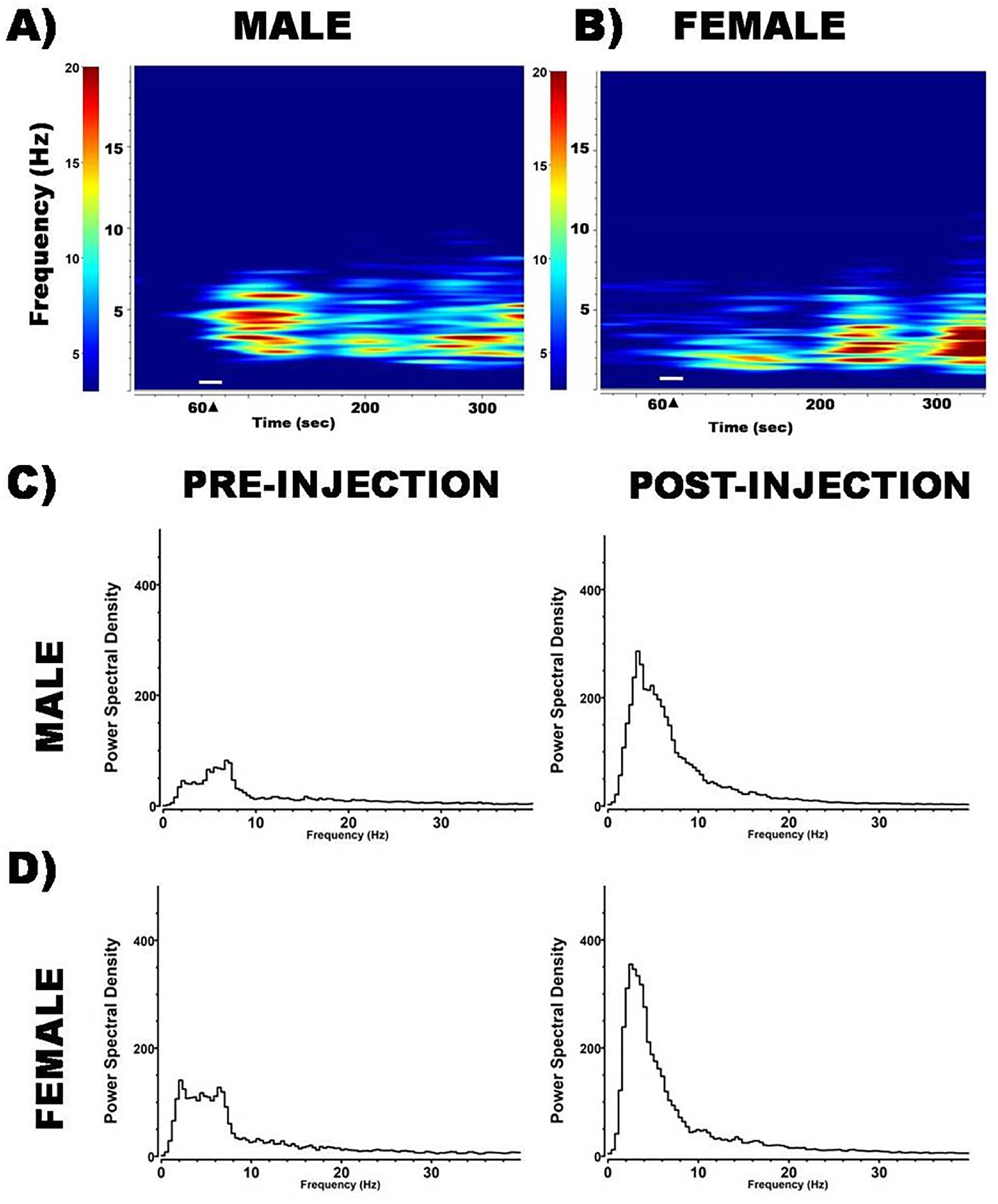
Similar effects of intra-LC clonidine on mPFC network activity. Time-frequency spectrograms of mPFC LFP activity from a male and female rat before and after (A and B) clonidine (50 ng in 200 nl). Power in different frequencies (ordinate) is indicated as color. The horizontal white line and arrowhead indicates the time of microinfusion. (C and D) Power spectral density plots corresponding to their respective time-frequency spectrograms shown above.

## DISCUSSION

The present results are consistent with previous findings of decreased MOR function in the LC of female compared to male rats (Guajardo *et al*, 2017). Decreased MOR mRNA expression and protein levels in the LC of female rats result in a decreased efficacy of DAMGO to inhibit LC neuronal activity. This sex difference was particularly apparent at doses (10 pg and 30 pg) that maximally inhibited male LC neurons. Notably, these sex differences in LC-MOR function translated to a sexually distinct impairment of performance of a prefrontal cortex-mediated cognitive task (Guajardo *et al*, 2017). The current demonstration that LC-MOR activation by a similar DAMGO (10 pg) dose selectively affects prefrontal cortical network activity of male rats links the previous molecular and behavioral findings. Taken together, the results suggest that decreased MOR expression in the LC of females relative to males, translates to an attenuation of the cortical response to LC-MOR activation, and this can account for sexually distinct cognitive consequences of LC-MOR activation. These sex-specific cognitive consequences are relevant for understanding sex differences in opioid abuse. Additionally, given the evidence that MOR activation in LC neurons may counteract stress effects on this system (Valentino *et al*, 2008), the present findings suggest that sex differences in MOR expression contribute to the increased vulnerability of females to the hyperarousal components of stress-related disorders (Gold and Chrousos, 2002; Koob, 1999; Wong *et al*, 2000).

### Relationship to previous studies

The LC regulates executive function and cognitive flexibility through its widespread projections to the prefrontal cortex (Loughlin *et al*, 1986; Swanson and Hartman, 1975). Previous studies demonstrated that selective pharmacological manipulations of LC neuronal activity are sufficient to affect cortical activity. For example, in anesthetized rats, compounds that inhibited LC discharge rate such as the α2-agonist, clonidine, shifted the power spectrum of cortical EEG towards a high-amplitude, low frequency state that was similar to that seen during slow wave sleep (Aston-Jones *et al*, 1981; Berridge *et al*, 1993). The effects of intra-LC DAMGO on cortical EEG activity recorded from screws inserted into the frontoparietal and fronto occipital regions showed a similar effect (Bagetta *et al*, 1990). Conversely, compounds that increased LC discharge rate such as the cholinergic agonist bethanechol, induced cortical EEG desynchronization characterized by low amplitude, high frequency activity (Berridge *et al*, 1991). Likewise, intra-LC microinfusion of CRF increases LC discharge rate, and produces cortical EEG desynchronization (Curtis *et al*, 1997). Notably, LC activation produced by CRF shifts the mode of LC discharge towards a high tonic state that is thought to facilitate behavioral flexibility (Curtis *et al*, 1997; Valentino and Foote, 1987, 1988). Consistent with this, relatively low doses of CRF injected into the LC improve cognitive flexibility in an attentional set-shifting task mediated by the mPFC, and increase c-fos (an indicator of neuronal activation) in mPFC neurons (Snyder *et al*, 2012). Taken together, these results suggest that alterations in LC neuronal activity are sufficient to alter mPFC activity and impact mPFC-mediated cognitive functions. Although MORs are highly expressed in LC neurons, and engaging MOR in the LC impairs a mPFC-mediated cognitive task, its effects on mPFC activity have not been documented previously. Importantly, decreased MOR function in the LC of females relative to males resulted in sexually distinct consequences in mPFC-mediated cognitive tasks, implying sex differences in the impact of activating LC-MOR on mPFC neuronal activity. Therefore, in this study we quantified and compared the effects of activating LC-MOR on mPFC neuronal activity between male and female rats.

### Effects of LC-MOR activation on prefrontal cortex activity

Field potential measurements provide an excellent tool for the exploration of network activity in mPFC. Previous studies of cortical network activity demonstrated that inhibition of LC discharge by clonidine induced high-amplitude, low frequency cortical oscillations (Berridge *et al*, 1993; de Sarro *et al*, 1988). Like clonidine, a DAMGO dose that completely inhibited LC activity, increased high-amplitude, low frequency oscillations in the mPFC. However, this effect was selective to male rats. The effects on the mPFC network activity were artifacts of injection, as these effects were not reproduced with intra-LC infusion of ACSF. The effects of DAMGO on mPFC network activity were also time-dependent in that they peaked at 10-15 min, and recovered by 45-60 min. Although for females there was a trend for an immediate effect of DAMGO on mPFC activity, this did not achieve statistical significance and was not apparent at later times after the injection. The lack of effect of intra-LC DAMGO (10 pg) in females is consistent with the reduced effect of the same dose on LC activity previously described, and attributed to decreased MOR expression in the LC (Guajardo *et al*, 2017). Finally, the observation that intra-LC clonidine produces comparable effects on mPFC activity in males and females supports the notion that the sex differences in the DAMGO response are due to differences at the level of MOR expression in the LC.

### Sex Differences in Behavioral/Cognitive Endpoints of MOR Activation in the LC

Microinfusion of DAMGO into the LC produced sex-specific effects on a strategy-shifting task thought to be mediated by the mPFC (Guajardo et al., 2017). For males, DAMGO significantly increased the time to complete the task, in part because it increased premature responding. Overall, male rats administered DAMGO made more errors during the task than females, particularly the number of regressive errors, which is indicative of an inability to acquire and maintain the new strategy. Consequently, sex-differences in LC-MOR expression that result in an enhanced ability of DAMGO to regulate mPFC activity, translated to impairments in learning to shift strategies selectively in males. Interestingly, female rats administered DAMGO into the LC committed more perseverative errors than male rats, which is indicative of an impaired ability to shift from a previously learned rule. Perseverative responding has been attributed to brain regions other than the prelimbic region, which was the site of recordings in the present study, including the infralimbic mPFC (Baran *et al*, 2010). This may explain why DAMGO in the female LC can have behavioral effects in the absence of electrophysiological effects in the prelimbic mPFC. A prior study using a place recognition task demonstrated that female rats were more sensitive to disruption of the infralimbic cortex as indicated by increased perseverative responding (Baran *et al*, 2010). Taken with the present results, this suggests that DAMGO in the LC of females may sufficiently affect activity in cortical regions, other than the prelimbic mPFC, that regulate other cognitive processes. In this way engaging MOR receptors in the LC can have sexually distinct effects on cognitive processing.

mPFC activity and strategy-shifting are physiological and behavioral endpoints of LC-MOR activation, respectively. Because for both endpoints, the same intra-LC dose of DAMGO was tested (Guajardo *et al*, 2017) and the experimental time-courses were comparable, the temporal relationship between cortical activity and behavior can be assessed. The time during which DAMGO had a maximal effect on cortical activity in male rats (10-15 min post injection) corresponded to performance of the simple discrimination stage of the task. DAMGO (10 pg) tended to enhance side discrimination performance when compared to vehicle control in males only, suggesting that mPFC synchronization may protect against distraction and is optimal for learning simple tasks. At the time point corresponding to strategy-shifting stage of the task (45 min), mPFC network activity in males partially recovered while effects on behavioral performance were apparent. This suggests an enduring effect on mPFC neuronal function that may not be expressed as changes in network activity. Alternatively, the discrepancy in the temporal correlation between physiological and behavioral endpoints may be a function of not recording both simultaneously in the same subject.

## CONCLUSION

Taken with our previous findings, the current findings demonstrate that the consequences of activating LC-MOR on mPFC activity and function are greatly diminished in females as a result of decreased MOR expression. MOR in the LC serves to counter stress-elicited excitation that is mediated by CRF and promotes recovery. The impact of reduced LC-MOR influence during stress coupled with increased CRF receptor signaling in females (Bangasser *et al*, 2010; Bangasser *et al*, 2013) would be predicted to result in a prolonged hyperactivity of the LC-NE system in response to stress and hyperarousal symptoms that characterize stress-related disorders in females (Gold *et al*, 2002; Koob, 1999; Wong *et al*, 2000).

Sex differences in LC-MOR function also have implications for sex differences in opioid abuse. In male rats, repeated social stress causes an imbalance between endogenous opioids, and CRF which favors opioid regulation (Chaijale et al, 2013). The increased opioid influence in the LC would bepredicted to promote premature responding, which is indicative of impulsive behavior (Pattij *et al*, 2009) in males only (Guajardo *et al*, 2017). Impulsivity is a key feature associated with opioid abuse (Baldacchino *et al*, 2015). These cellular alterations in LC after chronic stressors, coupled with the cognitive consequences of LC-MOR activation may predispose males to opiate abuse. The present study underscores how sex differences at the molecular level can translate to sex differences in network activities that govern behavior and cognition.

## Acknowledgements

We acknowledge the technical assistance of Xiao-Yan Zhang and Dr. Andre Cutis.

